# cGAS-STING dependent type I IFN protects against *Leptospira interrogans* renal colonization in mice

**DOI:** 10.1101/2025.06.02.657349

**Authors:** Suman Gupta, James Matsunaga, Bridget Ratitong, Sana Ismaeel, Diogo G. Valadares, A. Phillip West, Nagaraj Kerur, Christian Stehlik, Andrea Dorfleutner, Jargalsaikhan Dagvadorj, Jenifer Coburn, Andrea J. Wolf, Suzanne L. Cassel, David A. Haake, Fayyaz S. Sutterwala

**Author notes:** These authors contributed equally (Suzanne L. Cassel, David A. Haake, Fayyaz S. Sutterwala).

## Abstract

*Leptospira interrogans* is the major causative agent of leptospirosis. Humans, canines and livestock animals are susceptible to *Leptospira* species and can develop fulminant disease. Rodents serve as reservoir hosts in which the bacteria colonize the renal tubules and are excreted in the urine. The host immune response to *Leptospira* spp. remains poorly defined. We show that *L. interrogans* induces a robust type I interferon (IFN) response in human and murine macrophages that is dependent on the cytosolic dsDNA sensor Cyclic GMP-AMP Synthase (cGAS) and the Stimulator of IFN Genes (STING) signaling pathway. Further, we show that mice deficient in the IFNα/β receptor subunit 1 (IFNAR1) or STING had higher bacterial burdens and increased renal colonization following infection *in vivo* suggesting that cGAS-STING-driven type I IFN is required for the host defense against *L. interrogans*. These findings demonstrate the significance of cGAS-STING-dependent type I IFN signaling in mammalian innate immune responses to *L. interrogans*.

**Author Summary:** Leptospirosis is a globally distributed zoonotic disease caused by spirochetes belonging to the genus *Leptospira*. While humans, livestock, and dogs can develop severe or even fatal illness upon infection, rodents typically serve as asymptomatic reservoir hosts. A defining feature of the leptospiral life cycle is the ability of the pathogen to colonize the kidney in these reservoir host species, leading to prolonged urinary shedding and environmental dissemination. Despite the significant global burden of leptospirosis, the innate immune pathways that detect this pathogen and prevent renal colonization remain poorly understood. In this study we demonstrate that *L. interrogans* induces a robust type I IFN cytokine response from macrophages. The induction of this type I IFN response is dependent on sensing cytosolic DNA by the cGAS-STING pathway. Using *in vivo* mouse models of *L. interrogans* infection we further show that activation of this pathway is required to control bacterial burdens and reduce long-term kidney colonization. This study is the first to demonstrate a critical role for cGAS-STING and type I IFN in controlling *L. interrogans* infection.

## Introduction

Leptospirosis is caused by didermal spirochetes belonging to the genus *Leptospira* with *L. interrogans* being responsible for most pathogenic cases (1). All vertebrates are susceptible to infection, and the manifestation of symptoms varies extensively in different hosts (2). Cattle and canines experience mild to severe illnesses, indicating a heightened sensitivity to leptospirosis. In contrast, rodents such as mice and rats typically remain asymptomatic while being chronically colonized in their kidneys, serving as reservoir hosts for *Leptospira* spp. and shedding the bacteria in their urine for long periods of time (3–5). This process leads to soil and water contamination, perpetuating the enzootic cycle of the disease. Humans are incidental hosts with many infections remaining asymptomatic. However, humans can develop multiorgan dysfunction which can prove fatal in 10-20% of cases (6).

Leptospires are stealth pathogens that evade recognition by components of the host innate immune system. Leptospires have been shown to evade TLR5 activation due to the periplasmic positioning of their endoflagella (7). Additionally, they avoid detection by NOD1/NOD2 by tightly associating peptidoglycan with the outer membrane lipoprotein LipL21 which prevents the release of immunologically active peptidoglycan fragments (8). Despite these immune evasion strategies, leptospires also stimulate innate immune responses via other pathways. In both human and murine cells, leptospires are potent agonists of TLR2, primarily through their tri-acylated lipoproteins (9,10). *Leptospira* possess lipopolysaccharide (LPS) in their outer membrane, which serve as a virulence factor (11). Leptospiral LPS is also known to be recognized by TLR4 in mice (12). TLR2 and TLR4 recognition of leptospires has been identified as critical for murine resistance to acute leptospirosis (13).

The role of macrophages in the detection and clearance of leptospires remains incompletely understood. Depletion of peritoneal macrophages in C57BL/6J mice was shown to increase sensitivity to leptospirosis, resulting in an increased bacterial burden (14–17). In addition, macrophage depletion led to increased kidney colonization (16,17). Leptospires were generally classified as extracellular bacteria, although more recent studies have suggested the presence of leptospires within macrophages (18–20). Leptospires may either exist freely in the cytosol or reside within specific cellular compartments, such as endosomes, lysosomes, or phagosomes (19). The fate of internalized leptospires is unclear, but recent findings indicate that leptospires do not replicate within mouse macrophages and that viable internalized leptospires can exit these cells over time (21). The precise mechanisms by which macrophages detect leptospires and contribute to their control remains inadequately understood.

Intracellular innate immune pathways that sense the presence of cytosolic nucleic acids can play a critical role in the generation of type I interferon (IFN) responses. Cyclic GMP-AMP synthase (cGAS) is one such intracellular sensor that can bind to double-stranded DNA in the cytosol (22–26). Upon binding, cGAS generates 2’3’-cyclic guanosine monophosphate-adenosine monophosphate (cGAMP), which activates stimulator of interferon genes (STING). This activation results in the recruitment and subsequent phosphorylation of Tank-binding kinase-1 (TBK1) and IFN regulatory factor 3 (IRF3), followed by the production and secretion of IFNa and IFNý. Type I IFN signaling through the IFNα/β receptor (IFNAR) induces the transcription of interferon-stimulated genes (ISGs) (27,28). The cGAS-STING pathway was initially shown to play a role in anti-viral host response; it has since been shown to have roles in immune responses to a variety of bacterial pathogens including *Listeria monocytogenes, Mycobacterium tuberculosis, Klebsiella pneumoniae*, and *Borrelia burgdorferi* (23,25,26,29). Bacteria that produce cyclic dinucleotides can directly activate STING independently from cGAS (30–33). Whether *Leptospira* can activate the cGAS-STING pathway and the physiologic relevance of type I IFN production in leptospirosis is currently unknown.

Because pathogenic leptospires possess genes involved in cyclic dinucleotide synthesis (34,35) and leptospires can be internalized by macrophages, we asked if cGAS-STING-mediated type I IFN signaling was involved in the host immune responses against leptospiral infection. In this study, we demonstrate that *L. interrogans* induces type I IFN production and the transcription of ISGs in both mouse and human macrophages. The type I IFN response was found to be dependent on effective phagocytosis and degradation of leptospires in the macrophages. Further, we show that the type I IFN induction in *L. interrogans*-infected macrophages involves the cGAS-STING pathway. Finally, the absence of IFNAR or STING *in vivo* resulted in increased susceptibility to *L. interrogans* infection and increased kidney colonization, highlighting the importance of cGAS-STING-driven type I IFN in mammalian host defense against *L. interrogans*.

## Results

### *Leptospira* induces a type I IFN response following infection of macrophages

We first evaluated the ability of mouse macrophages to induce a type I IFN response following infection with *L. interrogans*. Consistent with prior reports we observed that bone marrow-derived macrophages (BMDMs) infected with *L. interrogans* serovar Copenhageni strain Fiocruz L1-130 at a multiplicity of infection (MOI) of 100 elicited the secretion of TNF, IL-10, and IL-6 (**Fig 1A-C**) (13,36). Additionally, robust secretion of IFNβ was also observed (**Fig 1D and S1A**). Tank-binding kinase 1 (TBK1) and IFN regulatory factor 3 (IRF3) play central roles in the initiation of type I IFN responses following signaling by the mitochondrial antiviral signaling protein (MAVS), STING, and TICAM1 (37). We found *L*. *interrogans* infection of BMDM resulted in the phosphorylation of TBK1 and IRF3 consistent with the observed production of IFNβ (**Fig 1E**). We also assessed the transcript levels of *Ifna* and *Ifnb* in BMDMs and observed upregulation of both transcripts by 4 hours post-infection (**Fig 1F, G**). Similar kinetics for IFN-β transcript upregulation were observed in PMA-differentiated human THP-1 cells (**Fig 1H**) suggesting *L. interrogans* can induce a type I IFN response in both mouse and human macrophages.

**Fig 1.**
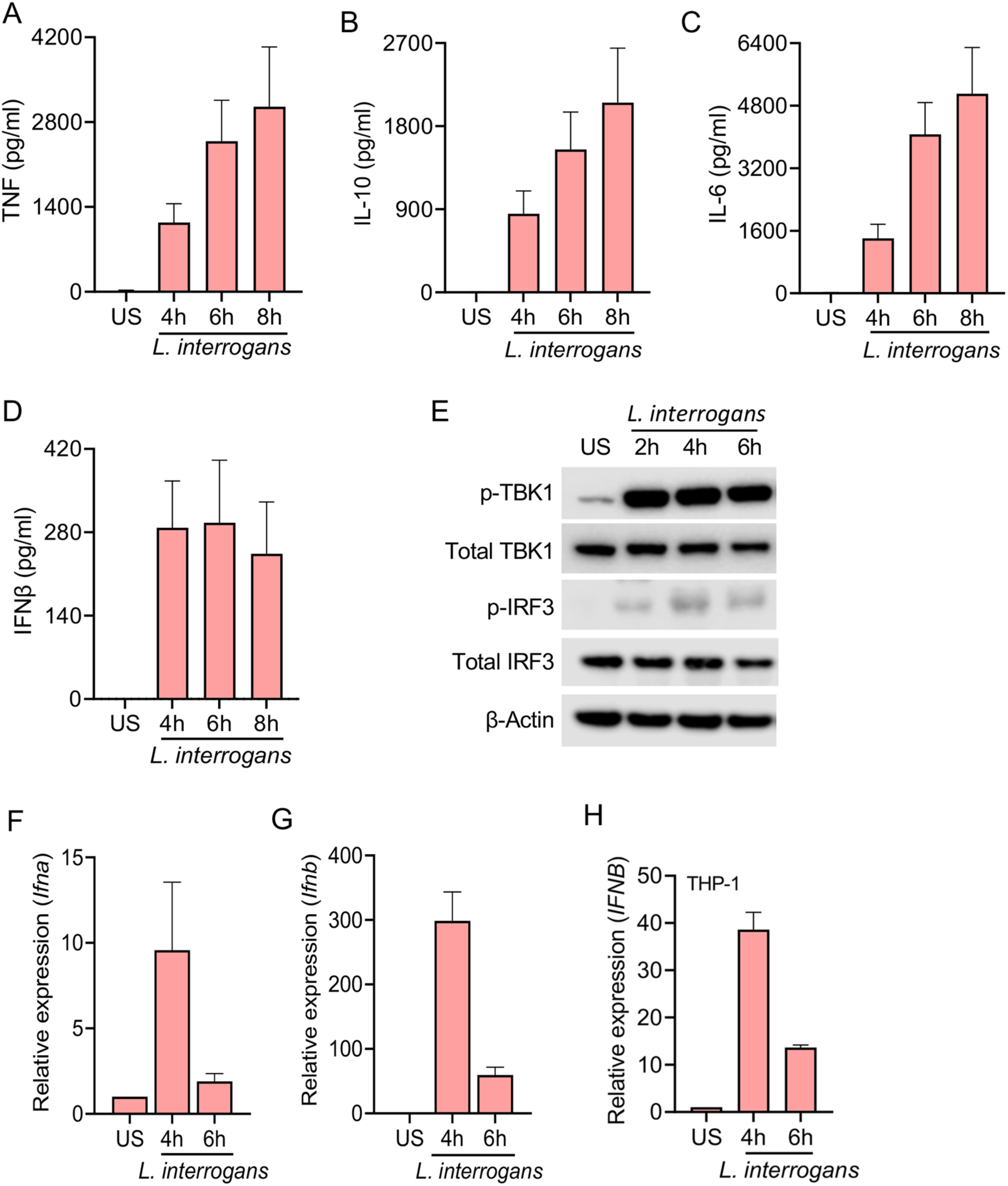
*L. interrogans* infection of macrophages induces a type I IFN response. (A-D) Quantification of TNF, IL-10, IL-6, and IFNβ by ELISA in culture supernatants of WT BMDMs infected with *L. interrogans* Fiocruz L1-130 (MOI 100) for 4, 6, and 8h. (E) Immunoblot for p-TBK1, total TBK1, p-IRF3, total IRF3 and β-actin in cell lysates of BMDMs infected with *L. interrogans* (MOI 100) for 2, 4, and 6h. Data are representative of three independent experiments. (F, G) Relative expression of *Ifna* and *Ifnb* transcript levels in BMDMs infected with *L. interrogans* at 4 and 6h post infection. (H) Relative expression of *IFNB* transcript levels in PMA-differentiated human THP-1 cells infected with *L. interrogans* at 4, 6, and 8h post infection. (A-D and F-H) Data are pooled from three independent experiments and expressed as the mean ± SEM.

### Phagocytic uptake of *L. interrogans* is required for effective IFNβ **secretion.**

To determine if *L. interrogans* were being phagocytosed, BMDMs were infected with pHrodo-labelled bacteria and increase in fluorescence dependent on phagosomal acidification was assessed. We found a robust increase in florescence of pHrodo-labelled bacteria with time suggesting bacteria were being taken up in acidic phagosomal compartment (**Fig 2A**). Additionally, wild-type (WT) BMDMs pretreated with cytochalasin D, an inhibitor of actin polymerization, prior to infection with pHrodo-labelled *L. interrogans* showed a reduction in fluorescence, suggesting that the uptake of bacteria into acidic subcellular compartments is via a phagocytic pathway (**Fig 2A**). After phagocytosis, bacteria-containing phagosomes recruit Rab5 to their surface, which is then replaced by LAMP1 upon lysosomal fusion (38–41). We found a subset of internalized bacteria colocalizing with Rab5 at 3 hours post infection (**Fig 2B**). Some bacteria were also found to be colocalizing with LAMP1 (**Fig 2B**) consistent with phagocytic uptake and processing of bacteria.

**Fig 2.**
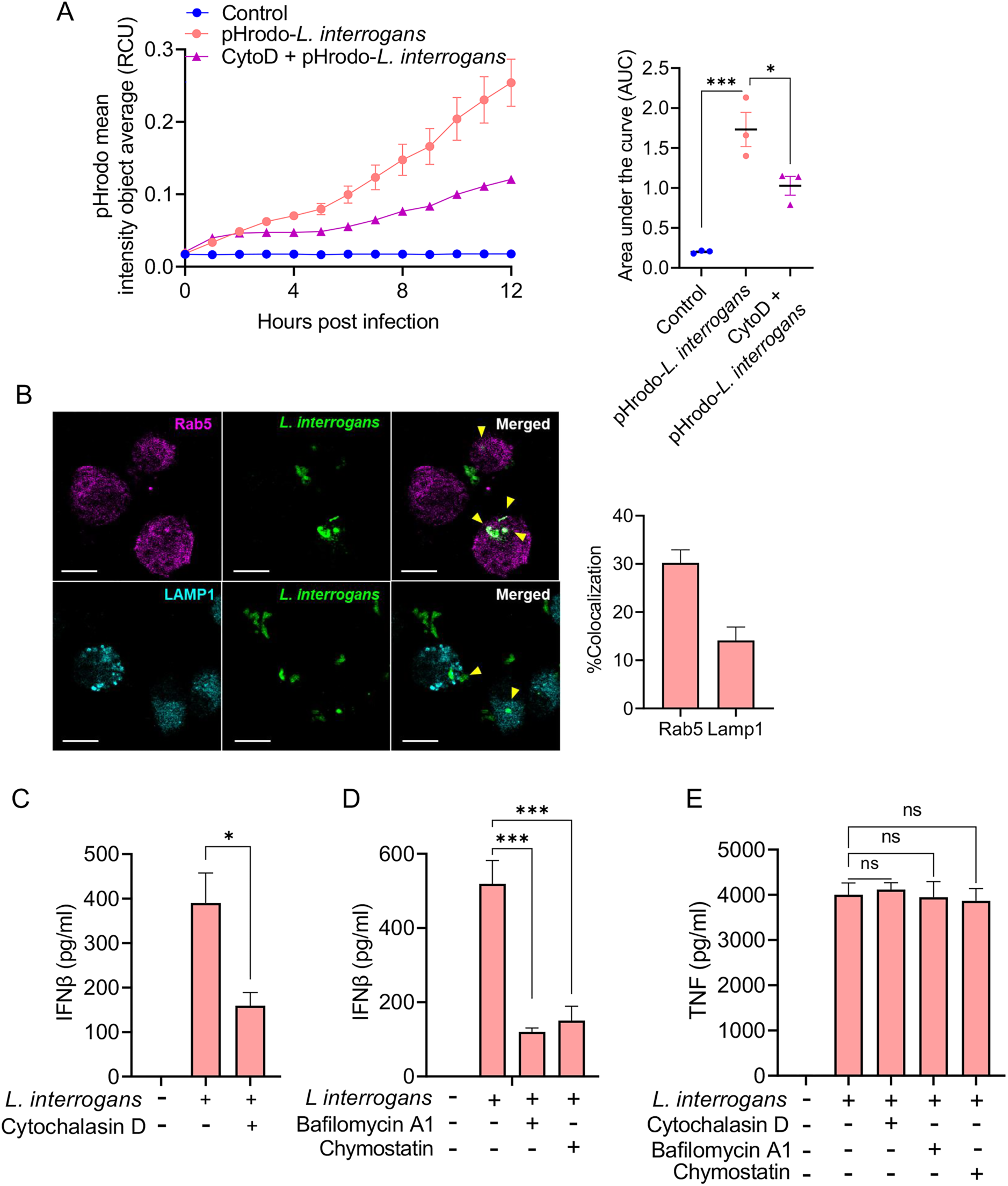
Phagocytosis of *L. interrogans* is required for macrophage driven IFNβ production. (A) Increase in mean fluorescence intensity per cell (Red Calibrated Unit; RCU) in WT BMDMs, after addition of *L. interrogans* Fiocruz L1-130 (MOI 100) labelled with pH-sensitive dye pHrodo, read by Incucyte SX5 reader at 1h intervals for a period of 12h in the presence or absence of cytochalasin D (2µM). Curves are representative of three independent experiments. Area under the curve (AUC) was calculated for the increase in pHrodo fluorescence intensity per cell in WT BMDMs in the presence or absence of cytochalasin D. Data are pooled from three independent experiments and expressed as the mean ± SEM. (B) Representative immunofluorescence image showing co-localization (arrow heads) of *L. interrogans* Fiocruz L1-130 with Rab5 and LAMP1 in BMDM at 3h post infection. Scale bar = 10 μm. Quantification of co-localization by pixel-pixel overlap calculated by JaCoP plugin in ImageJ. Data are pooled from three frames per experiment from three independent experiments. (C-E) Quantification of IFNβ and TNF by ELISA in culture supernatants of WT BMDMs pretreated with cytochalasin D, bafilomycin A1, and chymostatin, followed by infection with *L. interrogans* (MOI 100) for 6h. Data pooled from three experiments and expressed as the mean ± SEM. Statistical significance was calculated by two-way ANOVA; *p<0.05, ***p<0.001, ns=non-significant.

To determine if phagocytosis is required for the production of IFNβ, BMDMs were pretreated with cytochalasin D prior to infection with *L. interrogans*. Cytochalasin D treatment resulted in diminished IFNβ secretion (**Fig 2C**). Furthermore, treatment with bafilomycin A1, which inhibits phagosome and lysosomal acidification by blocking the vacuolar ATPase pump (42–44), and chymostatin, an inhibitor of phagosomal proteases (45,46), each resulted in reduced IFNβ following infection with *L. interrogans* (**Fig 2D**). Cytochalasin D, bafilomycin A1, and chymostatin did not affect TNF secretion from BMDM in response to infection with *L. interrogans* (**Fig 2E**). These findings suggest that *L. interrogans* phagocytosis by macrophages is crucial for a robust type I IFN response. Challenge of BMDM with heat-killed *L. interrogans* still resulted in IFNβ secretion indicating that live bacteria were not required for induction of the IFNβ response (**S1B Fig**). The induction of IFNβ secretion was also not dependent on the pathogenicity of the bacteria, as both pathogenic *L. interrogans* Fiocruz L1-130 and saprophytic *L. biflexa* serovar Patoc strain Patoc 1 strains induced similar levels of IFNβ (**S1C Fig**).

### Type I IFN signaling is required for effective control of *L. interrogans in mice in vivo*

Given that *Leptospira* induced a robust type I IFN response, we next assessed if the induction of type I IFN was important for the host’s ability to control bacterial replication in a mouse model of infection. C57BL/6J mice infected with *L. interrogans* do not develop fulminant disease; however, *L. interrogans* does chronically colonize the kidneys where they can be shed in the urine (47–49). To investigate the role of type I IFN signaling in acute *L. interrogans* infection, we used mice deficient in the IFNα/β receptor subunit 1 (IFNAR1). As expected, *in vitro* analysis of BMDMs from *Ifnar1*^-/-^ and WT mice showed no difference in IFNβ secretion following infection with *L. interrogans* (**Fig 3A**); however, the expression of ISGs, such as *Ifit1, Ifit3, Ifi44*, and *Zbp1* was significantly diminished in *L. interrogans* infected *Ifnar1*^-/-^ BMDMs compared to WT (**Fig 3B**).

**Fig 3.**
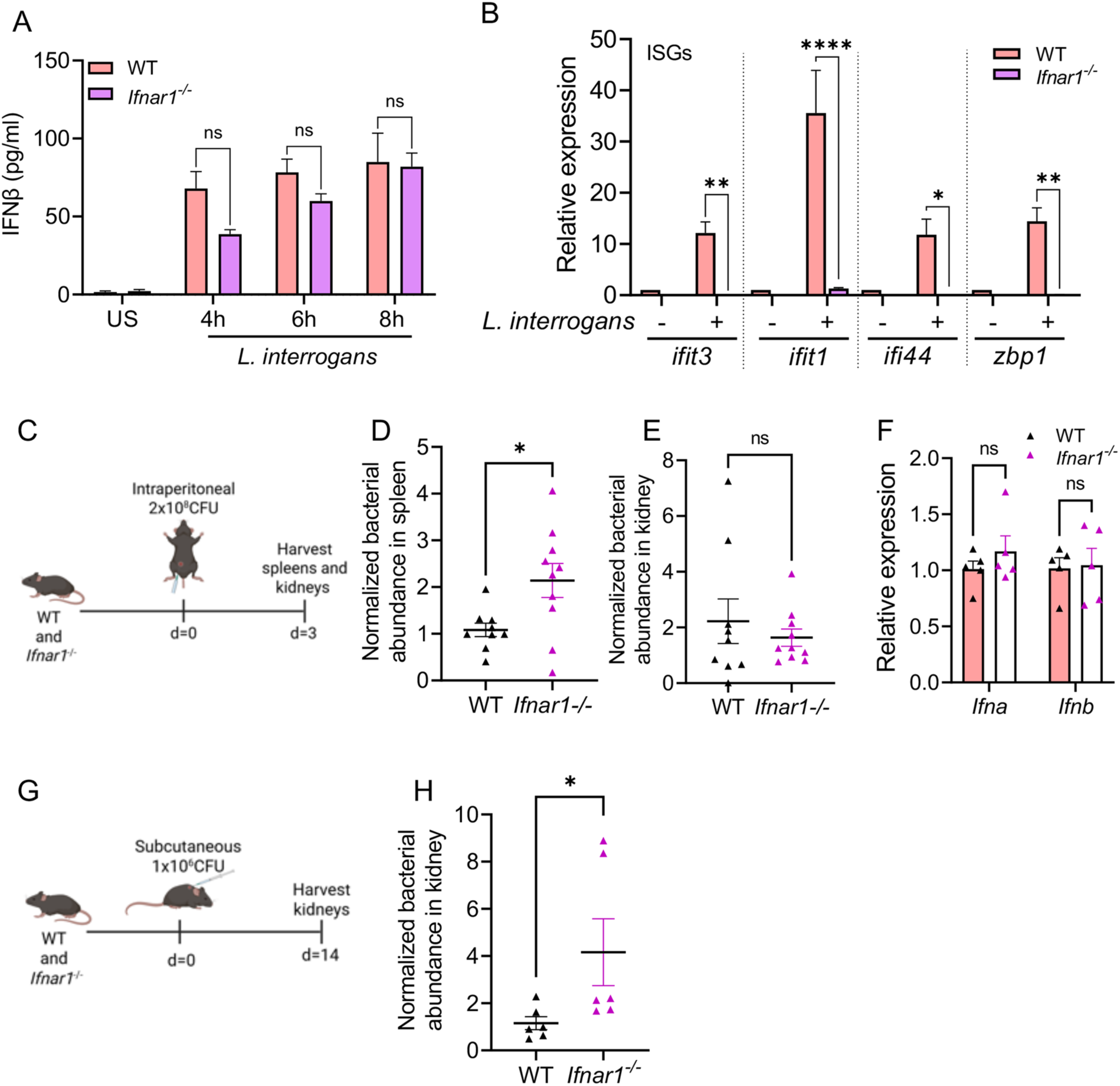
Increased susceptibility of IFNAR-deficient mice to *L. interrogans in vivo*. (A) Quantification of IFNβ by ELISA in culture supernatants of WT and *Ifnar1^-/-^* BMDMs infected with *L. interrogans* Fiocruz L1-130 (MOI 100) for 4, 6, and 8h. Data are pooled from three experiments and expressed as the mean ± SEM. Statistical significance was calculated by two-way ANOVA. (B) Relative expression of *ifit1, ifit3, ifi44,* and *zbp1* transcript levels in WT and *Ifnar1^-/-^* BMDMs infected with *L. interrogans* at 6h post infection. Data pooled from three experiments and expressed as the mean ± SEM. (C) Schematic of acute infection model; WT and *Ifnar1*^-/-^ mice were infected intraperitoneally with 2×10^8^ CFU of *L. interrogans* Fiocruz L1-130. (D, E) Bacterial abundance in the spleen and kidney at day 3 post-infection was assessed by qPCR for the *L. interrogans lipL32* gene and normalized to the eukaryotic *PPIA* gene. Data are pooled from two independent experiments: n=9 WT, n=10 *Ifnar1^-/-^*. (F) Relative expression of *Ifna* and *Ifnb* transcripts in spleen 24h post-acute infection in WT and *Ifnar1^-/-^* mice. n=5 WT, n=5 *Ifnar1^-/-^*. (G) Schematic of chronic infection model; WT and *Ifnar1^-/-^* mice were infected subcutaneously with 1×10^6^ CFU of *L. interrogans* Fiocruz L1-130. (H) Bacterial abundance in the kidney at day 14 post-infection was assessed by qPCR as above. Data are pooled from two independent experiments: n=6 WT, n=6 *Ifnar1^-/-^*. (D, E, H) Statistical significance was calculated by Mann-Whitney U test. (F) Statistical significance calculated by two-way ANOVA; *p<0.05, **p<0.01, ****p<0.0001, ns=non-significant.

Following an intraperitoneal *in vivo* infection with 2×10^8^ *L. interrogans* (**Fig 3C**), we found a higher bacterial burden in the spleen of *Ifnar1*^-/-^ mice compared to WT at 3 days post-infection (**Fig 3D**). In contrast, no difference in bacterial load was observed in the kidneys at 3 days post-infection (**Fig 3E**). As expected, transcript analysis of *Ifna* and *Ifnb* in the spleen at 24 hours post-infection revealed no differences between *Ifnar1*^-/-^ mice and WT mice (**Fig 3F**). Together these data demonstrate that *L. interrogans* induces the transcription of ISGs in *L. interrogans* infected macrophages and that type I IFN signaling through IFNAR1 is important for bacterial control in an acute leptospiral infection. To assess renal colonization by *L. interrogans*, mice were infected with 1×10^6^ spirochetes subcutaneously and kidney colonization assessed at 14 days post-infection (**Fig 3G**). *Ifnar1*^-/-^ mice had higher kidney colonization with *L. interrogans* compared to WT mice (**Fig 3H**). Taken together these data suggest that type I IFN signaling through IFNAR plays an important role *in vivo* in acutely controlling *L. interrogans* replication and limiting chronic kidney colonization.

### *L. interrogans*-induced type I IFN is cGAS-STING dependent

As the type I IFN response from BMDMs in response to *L. interrogans* was dependent on effective bacterial internalization and processing we hypothesized that a cytosolic pattern recognition receptor may be responsible for initiating this response. Additionally, the role of cytosolic DNA sensing in the innate immune response to *L. interrogans* remains unknown. To examine this, BMDMs from mice deficient in cGAS (*Cgas*^−/−^) and STING (*Sting1^gt/gt^*) were challenged with *L. interrogans* and *Ifna* and *Ifnb* transcript induction was assessed by qPCR. BMDM from *Cgas*^−/−^ and *Sting1^gt/gt^* mice showed a marked defect in the transcription of *Ifna* and *Ifnb* at 4 hours post-infection with *L. interrogans* compared to BMDMs from WT mice (**Fig 4A, B**). Consistent with this, BMDMs from *Cgas*^−/−^ and *Sting1^gt/gt^* mice also had diminished IFNβ secretion, but not TNF secretion, into culture supernatants as measured by ELISA compared to WT macrophages (**Fig 4C, D**). ISGs downstream of IFN signaling were also diminished in BMDMs from *Cgas*^−/−^ and *Sting1^gt/gt^* mice compared to WT following infection with *L. interrogans* (**Fig 4E**). A similar reduction in *IFNB* transcript levels was observed in the human monocytic THP-1 cell line with *CGAS* and *STING1* gene knockout (**Fig 4F**) indicating *L. interrogans* can activate the cGAS-STING pathway in both mouse and humans. Furthermore, leptospiral DNA directly transfected into BMDMs induced IFNβ production in a cGAS and STING dependent manner (**Fig 4G**).

**Fig 4.**
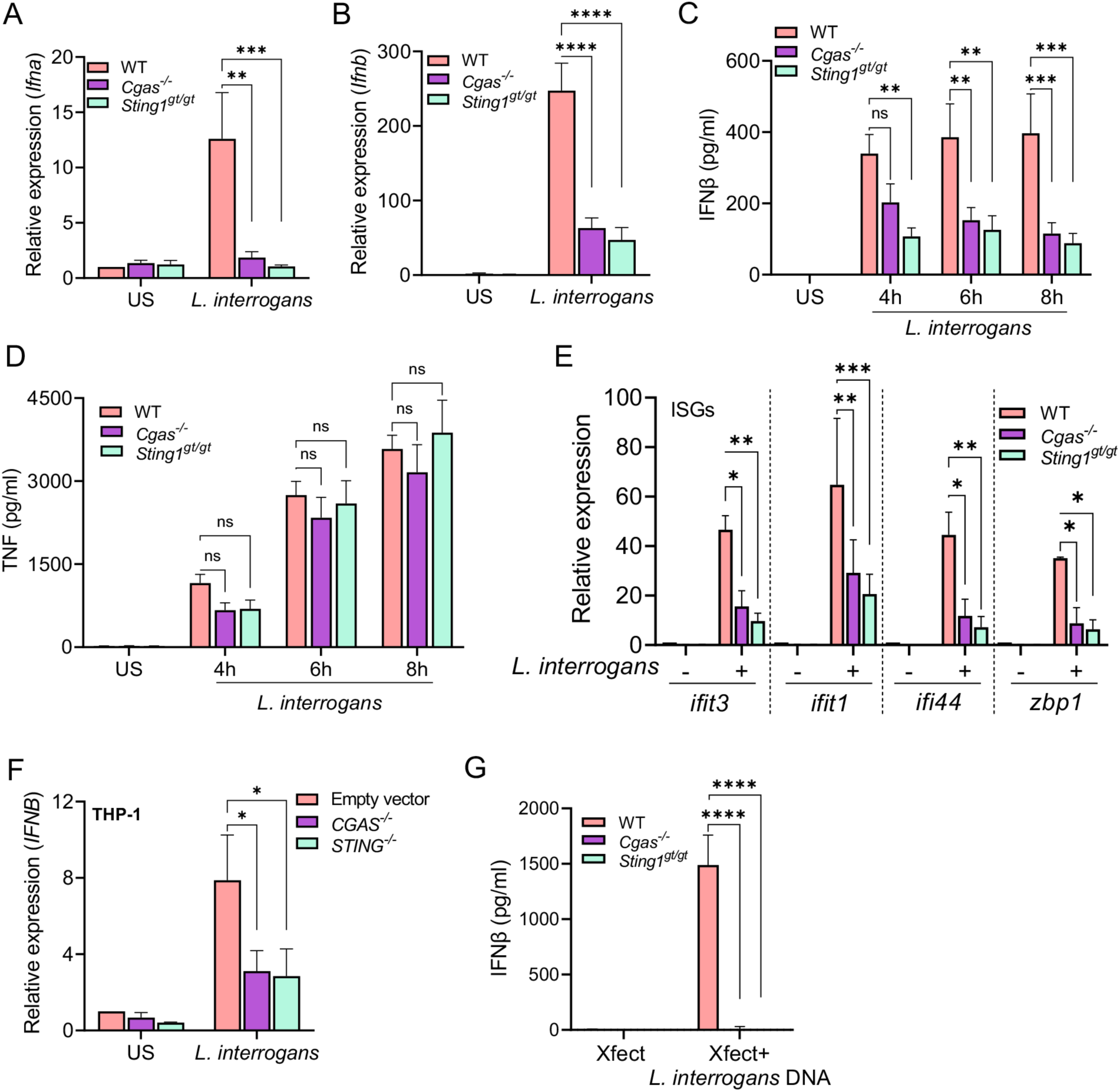
*L. interrogans* induced type I IFN is dependent on cGAS-STING. (A, B) Relative expression of *Ifna* and *Ifnb* transcripts in BMDMs from WT, *Cgas^-/-^*, and *Sting1^gt/gt^* mice 4h after infection with *L. interrogans* Fiocruz L1-130 (MOI 100). (C, D) Quantification of IFNβ and TNFα by ELISA in culture supernatants of WT, *Cgas^-/-^*, and *Sting1^gt/gt^* BMDMs infected with *L. interrogans* Fiocruz L1-130 (MOI 100) for 4, 6 and 8h. (E) Relative expression of *ifit1, ifit3, ifi44*, and *zbp1* transcript levels in BMDMs infected with *L. interrogans* (MOI 100) at 6h post-infection. (F) Relative expression of *IFNA* and *IFNB* transcript levels in PMA-differentiated human THP-1 cells infected with *L. interrogans* (MOI 100) at 4h post infection. (G) Quantification of IFNβ by ELISA in culture supernatants of WT, *Cgas^-/-^*, and *Sting1^gt/gt^* BMDMs transfected with 500 ng of *L. interrogans* DNA using Xfect transfection agent. (A-G) Data are pooled from three experiments and expressed as the mean ± SEM. Statistical significance was calculated by two-way ANOVA; *p<0.05, **p<0.01, ***p<0.001, ****p<0.0001, ns=non-significant.

To confirm that the differences in type I IFN responses seen in the absence of cGAS and STING is not due to differences in phagocytosis pathways we visualized the number of bacteria either bound or internalized by BMDMs. We found similar numbers of bacteria bound to or internalized by WT, *Cgas*^−/−^ and *Sting1^gt/gt^* BMDMs at 3 hours post-infection (**S2 Fig**). Taken together these data demonstrate that *L. interrogans* activates the cGAS-STING signaling cascade thereby inducing secretion of type I IFN and the subsequent transcription of ISGs in macrophages. Furthermore, as the production of type I IFN is cGAS dependent, these data suggest that cytosolic recognition of dsDNA initiates this pathway as opposed to bacterial derived cyclic dinucleotides.

### STING is required for effective control of *L. interrogans* in mice *in vivo*

To evaluate the contribution of STING in the control of *L. interrogans in vivo*, we intraperitoneally infected WT and *Sting1^gt/gt^* mice with *L. interrogans* and assessed bacterial burdens in the spleen and kidneys 3 days post-infection (**Fig 5A**). Higher bacterial burdens were observed in the spleen of *Sting1^gt/gt^* mice compared to WT (**Fig 5B**). No significant difference in bacterial burden was observed in the kidneys between WT and *Sting1^gt/gt^* mice (**Fig 5C**). Lower expression of *Ifna* and *Ifnb* was seen in the spleen of *Sting1^gt/gt^* mice compared to WT at 24 hours post-infection (**Fig 5D**). To assess renal colonization by *L. interrogans*, WT and *Sting1^gt/gt^*mice were subcutaneously infected with 1×10^6^ spirochetes and kidney colonization assessed at 14 days post-infection (**Fig 5E**). *Sting1^gt/gt^* mice had higher kidney colonization with *L. interrogans* compared to WT mice (**Fig 5F**). Taken together these data suggest that cytosolic recognition of *L. interrogans* activates the cGAS-STING pathway to control *L. interrogans* replication *in vivo* and limits chronic kidney colonization.

**Fig 5.**
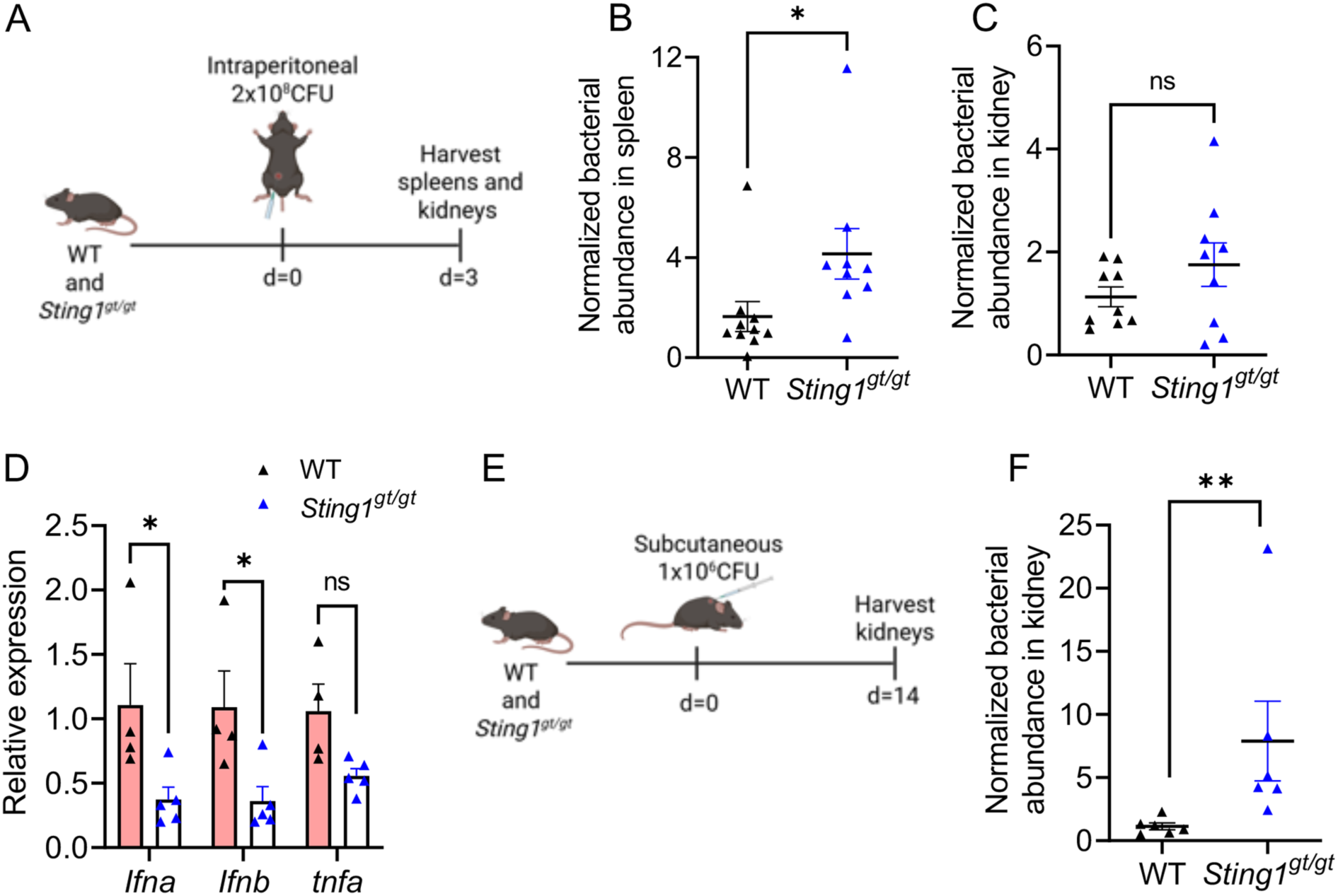
Increased susceptibility of STING-deficient mice to *L. interrogans in vivo*. (A) Schematic of acute infection model; WT and *Sting1^gt/gt^* mice were infected intraperitoneally with 2×10^8^ CFU of *L. interrogans* Fiocruz L1-130. (B, C) Bacterial abundance in the spleen and kidney at day 3 post-infection was assessed by qPCR for the *L. interrogans lipL32* gene and normalized to the eukaryotic *PPIA* gene. Data are pooled from two independent experiments: n=10 WT, n=9 *Sting1^gt/gt^*. (D) Relative expression of *Ifna, Ifnb*, and *tnf* transcripts in spleen 24h post-acute infection in WT and *Sting1^gt/gt^* mice. n=5 WT, n=5 *Sting1^gt/gt^*. Statistical significance was calculated by two-way ANOVA. (E) Schematic of chronic infection model; WT and *Sting1^gt/gt^* mice were infected subcutaneously with 1×10^6^ CFU of *L. interrogans* Fiocruz L1-130. (H) Bacterial abundance in the kidney at day 14 post-infection was assessed by qPCR as above. Data are pooled from two independent experiments: n=6 WT, n=6 *Sting1^gt/gt^*. (B, C, F) Statistical significance calculated by Mann-Whitney U test. *p<0.05, **p<0.01, ns=non-significant.

## Discussion

In this work, we have shown that *L. interrogans* induces a robust type I IFN response from human and mouse macrophages. The induction of type I IFN by *L. interrogans* was dependent on the cGAS-STING pathway as macrophages deficient in either cGAS or STING had markedly diminished expression of type I IFN and ISGs. Interestingly, the *L. interrogans* mediated production of type I IFN required bacterial phagocytosis and phagosomal acidification. We further demonstrate that the induction of type I IFN plays an important role in the host response to infection with *L. interrogans in vivo*; mice deficient in STING or IFNAR displayed increased susceptibility to *L. interrogans* infection and had increased kidney colonization at 14 days after subcutaneous challenge.

Leptospires are largely thought to be extracellular pathogens; however, studies have shown leptospires to be found within macrophages (19,21). The fate and impact of these intracellular leptospires has been unclear. Santecchia et al. demonstrated that *L. interrogans* can actively enter into both mouse and human macrophages (21). Surprisingly, they also found that the leptospires did not replicate within macrophages but could rapidly exit these cells intact (21). Using a pH-sensitive dye to label *L. interrogans*, we found that leptospires were being taken up in acidic sub-cellular compartments in macrophages. A fraction of internalized leptospires colocalized with Rab5 and LAMP1, suggesting the presence of leptospires in phagocytic vesicles. We further found that the pharmacologic inhibitors cytochalasin D, bafilomycin A and chymostatin, that target phagocytic uptake, phagosomal acidification, and proteolytic activity, respectively, inhibited *L. interrogans* induced IFNβ production. These data suggest that the phagocytosis and phagosomal processing of *L. interrogans* is required for the activation of the cGAS-STING pathway. It remains unclear to what extent these internalized leptospires are killed or damaged resulting in bacterial DNA release into the macrophage cytosol.

The cGAS-STING pathway can drive the production of type I IFN in response to pathogen-derived DNA as well as endogenous host DNA (50–53). cGAS can bind to dsDNA and produce cGAMP, which then binds to and activates STING on the endoplasmic reticulum membrane (51,53–55). This results in TBK1 phosphorylation, which phosphorylates the transcription factor IRF3 and drives the induction of type I IFN (56–58). We observed that *Leptospira* induces a type I IFN response, coinciding with the phosphorylation of TBK1 and IRF3, supporting a role for cGAS-STING in the detection of intracellular *L. interrogans*. Several intracellular bacterial pathogens, including *Salmonella enterica* serovar Typhimurium, *L. monocytogenes* and *M. tuberculosis* have been shown to activate the cGAS-STING pathway to drive type I IFN production (23,25,59). Extracellular bacteria, including *Klebsiella pneumoniae* and *Pseudomonas aeruginosa*, have also been shown to activate the cGAS-STING pathway (26). The spirochete *B. burgdorferi* can activate the cGAS-STING pathway in macrophages and fibroblasts (29). Although cGAS-STING did not appear to play a role in control of *B. burgdorferi* burdens *in vivo*, inflammation and joint pathology were diminished in cGAS-deficient mice infected with *B. burgdorferi* (29). The mechanism by which cGAS-STING recognizes *L. interrogans* remains unclear. Bacterial derived cyclic dinucleotides have been shown to directly activate STING (30–32). Although *L. interrogans* can produce cyclic dinucleotides (34,35), our finding that cGAS is required for *L. interrogans*-driven type I IFN production suggests that sensing of cytosolic DNA is required to engage the cGAS-STING pathway in response to *L. interrogans* and that *L. interrogans* derived cyclic dinucleotides do not directly activate STING. As internalization and phagocytic processing of *L. interrogans* is required for type I IFN production by macrophages it is possible that internalized spirochetes either shed DNA or are damaged within the phagosome resulting in release of bacterial DNA that can be recognized by cytosolic cGAS. Another possibility is that infection of macrophages with *L. interrogans* results in mitochondrial stress resulting in the release of mtDNA which can then activate cGAS-STING (59). It should be noted that other pattern recognition receptors, including RIG-I/MDA5/MAVS, TLR3, TLR4, TLR7, TLR8, and TLR9, may be involved in the induction of type I IFN responses (60–67). The role of these pattern recognition receptors in driving the type I IFN response to *L. interrogans* will require further investigation.

The murine model of leptospirosis remains poorly characterized, leaving many questions unanswered regarding the host response to *Leptospira* spp. A recent study by Papadopoulos et al. on the progression of acute leptospirosis in mouse models provides valuable insights into how severe leptospirosis in mice mimics aspects of human disease (68). This study attributes the observed mouse mortality in the acute leptospirosis model to severe myocarditis and neutrophil infiltration, in the absence of a significant cytokine storm (68). Our *in vivo* infection of *Sting1^gt/gt^*and *Ifnar1^-/-^* mice with *L. interrogans* revealed a role for type I IFN in both the early control of bacterial replication as well as in chronic kidney colonization with *L. interrogans*. The effect of the *L. interrogans*-driven type I IFN response on neutrophil function in vivo remains to be evaluated. In addition, the contribution of other cell types, such as renal tubular epithelial cells, in the production of type I IFN in response to *L. interrogans* will be important to determine.

In conclusion, our study is the first to show that *L. interrogans* activates the cytosolic cGAS-STING DNA sensing pathway in mouse and human macrophages resulting in a type I IFN response. Future studies are required to determine the source of cytosolic DNA that is recognized by cGAS-STING following *L. interrogans* infection of macrophages. Additionally, determining the intracellular fate of *L. interrogans* will be important in understanding if bacterial damage and release of bacterial DNA is a prerequisite for activation of cGAS-STING. We also found that activation of STING and signaling through IFNAR are required for controlling *L. interrogans* replication and kidney colonization in mice in vivo. Evaluating the role of type I IFN production during the course of human disease remains an unresolved question and may provide insights into species specific responses.

## Materials and methods

### Ethics statement

All studies with mice were performed in accordance with the recommendations in the Guide for the Care and Use of Laboratory Animals of the National Institutes of Health and were reviewed and approved by the Institutional Animal Care committee at Cedars-Sinai Medical Center (CSMC IACUC #006777).

### Mice

Wild-type (WT) C57BL/6J (JAX stock# 000664) mice were purchased from Jackson laboratories and used as controls unless otherwise stated. The generation of *Sting1^gt/gt^* (JAX stock# 017537), *Cgas*^-/-^ (JAX stock# 026554) and *Ifnar1*^-/-^ (JAX stock# 028288) have been previously described (69–71). Mice were bred and maintained in a specific-pathogen free facility. Both male and female mice 6–12 wk of age were used; however, mice were sex and age matched for individual experiments.

### Bacterial culture

*Leptospira interrogans* serovar Copenhageni strain Fiocruz L1-130 and *Leptospira biflexa* serovar Patoc strain Patoc 1 were grown in Hornsby-Alt-Nally (HAN) media (72). Cultures were incubated at 30°C and passaged weekly. *L. interrogans* was passaged a maximum of five times after collection from kidneys of infected Golden Syrian hamsters. Culture densities were determined by diluting 10 μL of culture into phosphate-buffered saline (PBS) and counting by darkfield microscopy with a Zeiss Axio Lab A1 or AmScope B340 Series microscope. Prior to experiments, leptospires were centrifuged at 9,000 x g for 4 min at room temperature. For *in vitro* experiments, leptospires were resuspended in DMEM to a final density of 1 x 10^9^/mL. For heat killing, the leptospires were heated at 56°C for 30 min. For labelling of bacteria with pHrodo™ Red, succinimidyl ester (Invitrogen # P36600), leptospires were incubated for 45 min at room temperature with 0.1 mM dye. Labelled bacteria were washed and then incubated with BMDMs to analyze increase in fluorescence using an Incucyte SX5 Live-Cell Analysis System (Sartorius). For *in vivo* experiments, leptospires were resuspended in HAN media to a density of 1 x 10^9^/mL following centrifugation of the culture. After resuspension, 200 μL was used to inoculate mice for acute infection; for renal colonization, the cell suspension was diluted 100-fold prior to inoculation of mice with 100 μL.

### *In vitro* stimulation of bone marrow–derived macrophages

Bone marrow–derived macrophages (BMDMs) were generated from cells collected by flushing mouse femurs and tibias and culturing the cells in 10 mm dish for 6-7 days in DMEM (Corning #10-013-CV) with 10% FBS (R&D systems #S12450H), penicillin-streptomycin (Gibco # 15140122), and L-glutamine (Gibco #25030-081) in the presence of 20% L929 cell-conditioned medium. Upon differentiation, BMDMs were harvested using Versene (Gibco# 15040066) and used for subsequent experiments. BMDMs were either left unstimulated or infected with *L. interrogans* Fiocruz L1-130 or *L. biflexa* Patoc I at multiplicity of infection (MOI) of 100 for 4, 6, and 8 h. Wherever indicated, BMDMs were pre-incubated for 45-60 mins with 2µM cytochalasin D (Sigma-Aldrich #C8273), 400nM bafilomycin A (Invivogen #tlrl-baf1) and 100µM chymostatin (Caymen Chemical #15114) before addition of bacteria. For transfection of leptospiral DNA, 500 ng of DNA isolated from *L. interrogans* Fiocruz L1-130 cultures was incubated with Xfect transfection reagent (Takara Bio #631317) for 10 min and BMDMs were treated with the mixture for 6 h. After respective stimulations, supernatants were collected for quantification of cytokines and cells were preserved in RNAlater Stabilization Solution (Invitrogen #AM7020) for RNA isolation using RNeasy Mini kit (Qiagen #74106) as per the manufacturers’ instruction. Cytokines and chemokines were quantified in the cell culture supernatants using mouse ELISA kit for TNF (Invitrogen #88-7324-88), IFNβ (R&D Systems #DY8234), IL-10 (R&D Systems #DY417) and IL-6 (R&D Systems #DY406) following the manufacturers’ protocol.

### Immunofluorescence staining and confocal microscopy

3×10^4^ BMDMs were seeded in an 8-well Millicell EZ Slide (Millipore Sigma #PEZGS0816) and incubated for 3 h with *L. interrogans* Fiocruz L1-130 at MOI 100 or left unstimulated. Cells were washed with PBS three times and fixed with 4% PFA (Electron Microscopy Sciences #15710) for 10 minutes at room temperature. For analyzing bound bacteria, after fixation, cells were not permeabilized and proceeded with immunofluorescence (IF) staining. For analyzing total bound and internalized bacteria, fixed cells were permeabilized using 0.1% TritonX-100 (VWR Life Science # 9002-93-1) for 10 min. Cells were then incubated in IF blocking buffer containing 2.5% BSA (RPI # A30075) and normal donkey serum (Abcam #ab7475) in PBS (Corning #21-040-CV) for 1 h at room temperature. Primary antibodies: rat anti-mouse LAMP1 (clone 1D4B, eBioscience), mouse anti-mouse Rab5 (clone rab5-65, Sigma-Aldrich), and rabbit anti-sera raised against *L. interrogans* Fiocruz L1-130 were diluted in IF blocking buffer and added to cells for overnight incubation at 4°C. Cells were washed three times with PBS and AF488 anti-rabbit IgG, AF555 anti-mouse IgG, and AF647 anti-rat IgG diluted in IF buffer were incubated with the cells for 1 h at room temperature. Cells were washed three times with PBS before removing the well borders. VECTASHIELD Vibrance Antifade Mounting Medium with DAPI (Vector Laboratories #H-1800) was added on top of each well of stained cells and sealed with a coverslip. Imaging was done using either Zeiss Z780 confocal microscope or Leica Stellaris confocal microscope. For counting the number of bacterial puncta that are bound vs the total number of bacteria bound and internalized, the Image J manual cell counting tool was used. For analyzing the percentage of total bacteria that colocalizes with Rab5 or LAMP1, the JaCoP pixel-based colocalization tool was used to calculate the Manders’ coefficient using manual thresholding.

### *In vivo* infections

For acute infection, mice were injected intraperitoneally with 2×10^8^ bacteria and on day 3 following infection spleen and kidney were harvested and stored in RNAlater Stabilization Solution (Invitrogen #AM7020) for further processing. Tissues were weighed and homogenized in PBS with 2.8 mm ceramic beads (Omni #19-646) using a FastPrep-24 Classic bead beating grinder and lysis system (MP # 116004500). Subsequently DNA was extracted from appropriate tissue amount using DNeasy Blood and Tissue kit (Qiagen # 69506) using the manufacturers’ protocol. To assess kidney colonization, mice were injected subcutaneously with 1×10^6^ bacteria and on day 14 following infection the kidney was harvested and processed as above.

### qPCR

cDNA was synthesized from the isolated RNA using PrimeScript RT master mix (Takara Bio #RR036A) as per the manufacturers’ protocol. Transcripts were amplified with the primers listed in **S1 Table** using PowerUP SYBR Green PCR master mix (Applied Biosystems #25742) in Applied biosystems ViiA7 96 well system. The transcript levels were normalized to housekeeping gene β-actin and data are shown as fold change of transcripts relative to mean of WT group using the ΔΔ threshold cycle (Ct) method. For quantifying the relative abundance of bacteria in kidneys and spleen of infected mice, the DNA was amplified with primers specific to the *lipL32* gene for *L. interrogans* and normalized to eukaryotic tissue inputs by amplification of *PPIA* housekeeping gene. Data are shown as relative abundance of bacteria normalized to tissue inputs and compared to the WT group on day 3 post infection using the ΔΔ Ct method.

### Cell lines

Human monocyte cell line THP-1 with *CGAS* and *STING1* gene knockout were generated by CRISPR/Cas9 as previously described (73). THP-1 monocytes were grown in RPMI-1640 medium (ATCC #30-2001) with 10% FBS (R&D systems #S12450H), penicillin-streptomycin (Gibco # 15140122), L-glutamine (Gibco #25030-081) and β-mercaptoethanol (Thermo Fisher # 21985023). THP-1 monocytes were differentiated using 50 nM PMA for 3-4 h, followed by an overnight rest period. The cells were then infected with *L. interrogans* Fiocruz L1-130 at MOI 100; 4 h later the cells were harvested for RNA isolation.

### Western blotting

Cell lysates were prepared in radioimmunoprecipitation assay (RIPA) lysis buffer (Cell Signaling Technology #9806S) with Halt Protease and Phosphatase Inhibitor Cocktail (Thermo scientific #78440). Proteins were separated on NuPAGE Bis-Tris Mini Protein Gels, 4–12%, 1.0–1.5 mm and transferred to a polyvinylidene difluoride (PVDF) membrane using either the XCell II blotting system (Invitrogen #EI0002) or iBlot 2 Transfer system (Invitrogen #IB24002). Membranes were blocked with 1% BSA in tris-buffered saline (TBS) with 0.1% Tween-20 and incubated with primary antibody overnight at 4°C. Primary antibodies from Cell signaling technologies (CST) were used as follows: TBK1 (CST #3504S), p-TBK1 (CST #5483S), IRF3 (CST #4302S), p-IRF3 (CST #4947S) and β-actin (CST #4970S). After washing, membranes were incubated with HRP-tagged anti-rabbit IgG (Cell Signaling Technology #7074S). Membranes were developed using SuperSignal West Pico PLUS (Thermo Scientific #34580) or Femto substrate (Thermo Scientific #34094) and imaged in LI-COR Odyssey Fc.

### Statistics

Data were graphed and the indicated statistical tests performed using GraphPad Prism 9 software. If not otherwise stated, analysis was performed with Unpaired *t* test with Welch’s correction for single comparison or Two-way ANOVA for multiple comparisons.

## Acknowledgments

This work was supported by National Institutes of Health grants P01 AI168148 (D.A.H.), R01 AI175101 (F.S.S. and J.D.), and R01 AI177233 (F.S.S. and S.L.C.).

## Author contributions

Conceptualization, S.G., S.L.C., F.S.S. and D.A.H.; Methodology, S.G., J.M., J.C., F.S.S. and D.A.H.; Investigation, S.G., J.M. and B.R.; Writing – Original Draft, S.G., F.S.S. S.L.C. and D.A.H.; Writing – Review & Editing, S.G., F.S.S. S.L.C. and D.A.H.; Funding Acquisition, S.L.C., J.D., F.S.S. and D.A.H.; Resources, A.P.W., N.K., C.S. and A.D.; Supervision, S.L.C., F.S.S. and D.A.H.

## Figure Legends

**S1 Fig.**
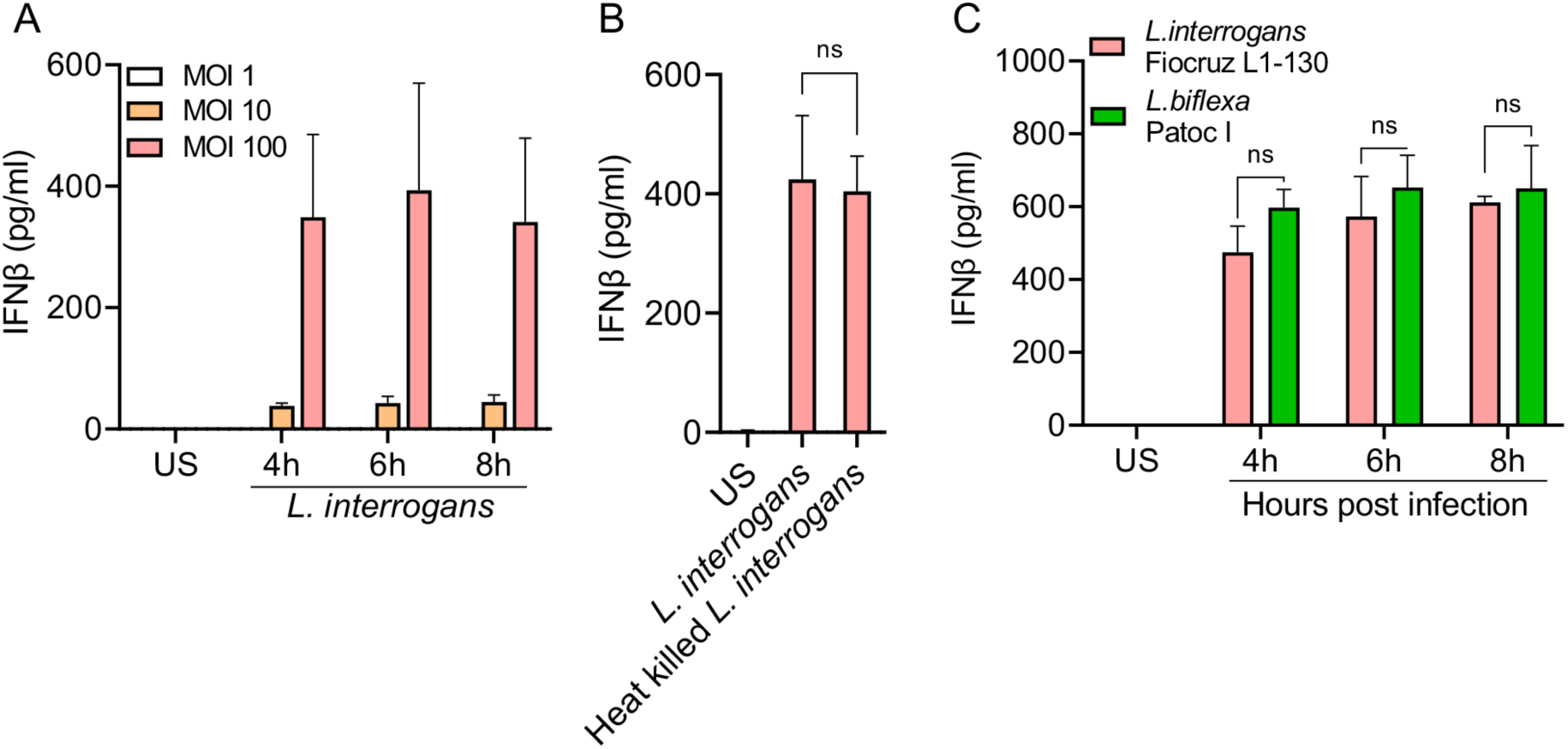
Analysis of IFNβ production from BMDM infected with *Leptospira.* (A) Dose dependent induction of IFNβ was quantified by ELISA in culture supernatants of WT BMDMs infected with *L. interrogans* (MOI 1, 10, 100) for 4, 6, and 8h. Data are pooled from two independent experiments and expressed as the mean ± SD. (B) Quantification of IFNβ levels by ELISA in culture supernatants of WT BMDM infected with live or heat-killed *L. interrogans* (MOI 100) at 6h post-infection. Data are pooled from three independent experiments. Statistical significance was calculated by unpaired Student’s *t* test. (C) Quantification of IFNβ levels by ELISA in culture supernatants of WT BMDM infected with *L. interrogans* Fiocruz L1-130 (MOI 100) or *L. biflexa* Patoc 1 (MOI 100) at 4, 6, and 8h post-infection. Statistical significance calculated by two-way ANOVA. ns=non-significant.

**S2 Fig.**
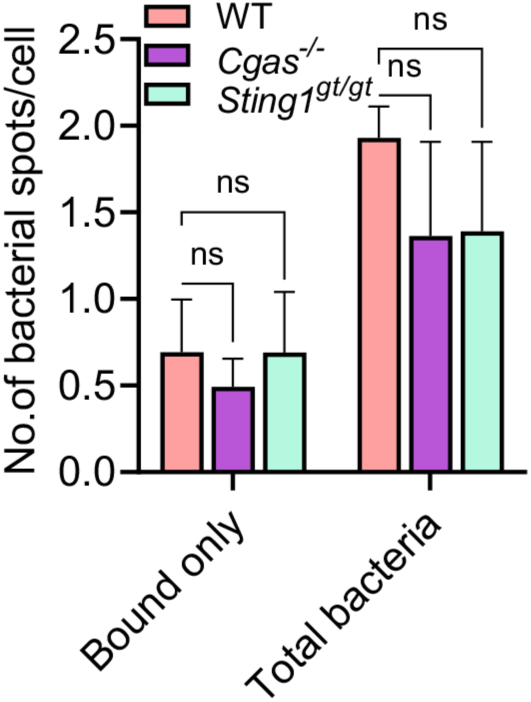
Analysis of *L. interrogans* binding and internalization by BMDM. Quantification of immunofluorescence images of bacteria bound or total bacteria (bound and internalized) in WT, *Cgas^-/-^* and *Sting1^gt/gt^* BMDMs at 3h post infection with *L. interrogans* (MOI 100). Data are pooled from three frames per experiment from three independent experiments. Statistical significance was calculated by two-way ANOVA. ns=non-significant.

**S1 Table:**
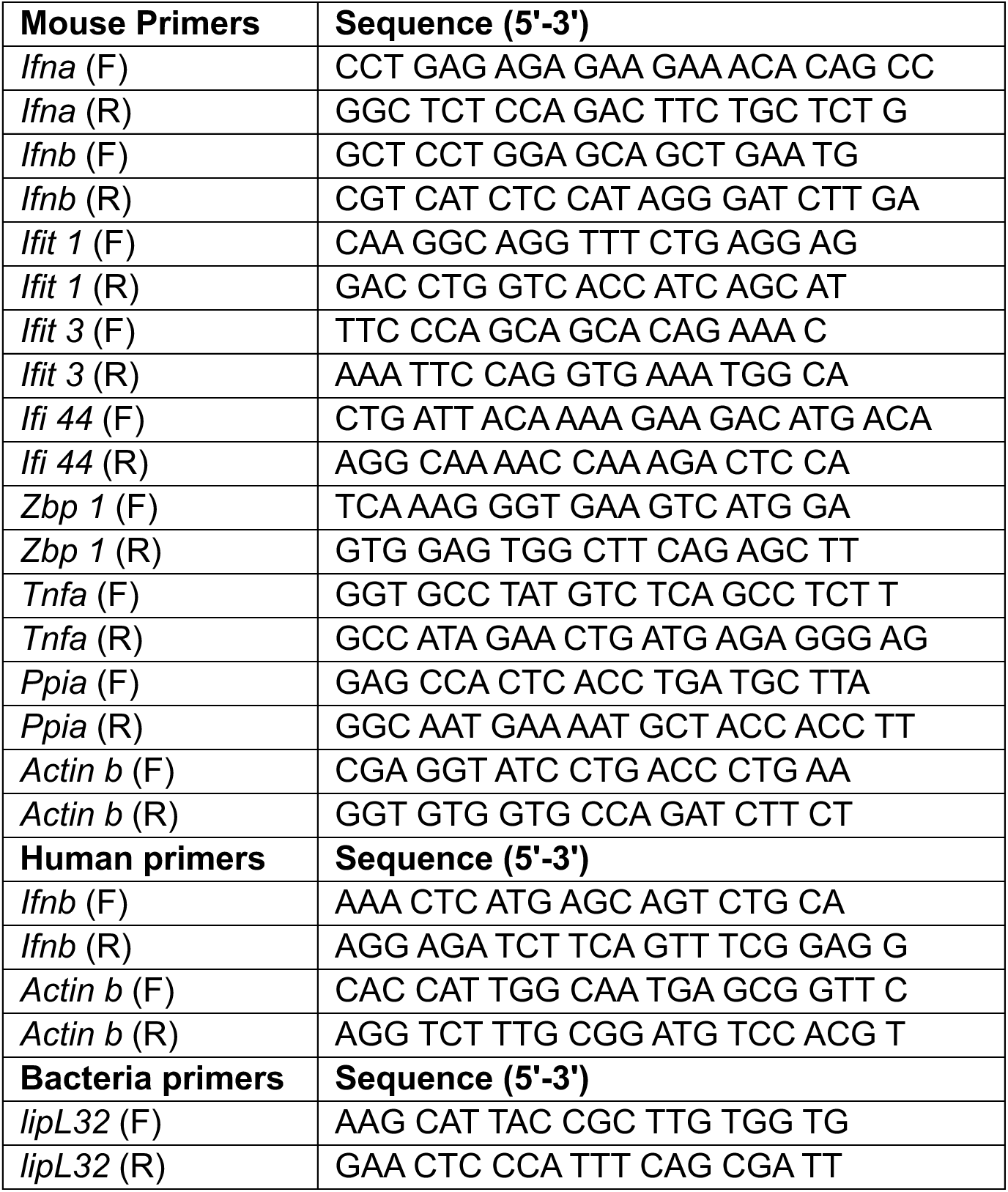
Primers used for qPCR.

